# Assessment of ACE2, CXCL10 and Their co-expressed Genes: An In-silico Approach to Evaluate the Susceptibility and Fatality of Lung Cancer Patients towards COVID-19 Infection

**DOI:** 10.1101/2020.05.27.119610

**Authors:** Tousif Bin Mahmood, Afrin Sultana Chowdhury, Mehedee Hasan, Md. Mezbah-Ul-Islam Aakil, Mohammad Imran Hossan

**Affiliations:** Department of Biotechnology and Genetic Engineering, Noakhali Science and Technology University, Noakhali-3814, Bangladesh

**Keywords:** COVID-19, SARS CoV-2, Lung Cancer, LUAD, LUSC, ACE2, CXCL10

## Abstract

**Background:** COVID-19 is a recent pandemic that started to spread out worldwide from Wuhan, China. This disease is caused by a newly discovered strain of the coronavirus, namely SARS CoV-2. Lung cancer patients are reported to be more susceptible to COVID-19 infection. To evaluate the probable reasons behind the excessive susceptibility and fatality of lung cancer patients to COVID-19 infection, we targeted two most crucial biomarkers of COVID-19, ACE2 and CXCL10. ACE2 plays a vital role in the SARS CoV-2 entry into the host cell while CXCL10 is a cytokine mainly responsible for the lung cell damage involving in a cytokine storm.

**Methods:** Firstly, we used the TIMER, UALCAN and GEPIA2 databases to analyze the expression and correlation of ACE2 and CXCL10 in LUAD and LUSC. After that, using the cBioPortal database, we performed an analytical study to determine the genetic changes in ACE2 and CXCL10 protein sequences that are responsible for lung cancer development. Finally, we analyzed different functional approaches of ACE2, CXCL10 and their co-expressed genes associated with lung cancer and COVID-19 development by using the PANTHER database.

**Results:** Initially, we observed that ACE2 and CXCL10 are mostly overexpressed in LUAD and LUSC. We also found the functional significance of ACE2 and CXCL10 in lung cancer development by determining the genetic alteration frequency in their amino acid sequences. Lastly, by doing the functional assessment of the targeted genes, we identified that ACE2 and CXCL10 along with their commonly co-expressed genes are respectively involved in the binding activity and immune responses in case of lung cancer and COVID-19 infection.

**Conclusions:** Finally, on the basis of this systemic analysis, we came to the conclusion that ACE2 and CXCL10 are possible biomarkers responsible for the higher susceptibility and fatality of lung cancer patients towards the COVID-19.

## 1. Introduction

In recent times, the corona viral disease 2019 (COVID-19) has noted as the most alarming disease that occurs due to the infection of severe acute respiratory syndrome coronavirus 2 (SARS-CoV-2). This new strain is declared as the seventh modification of corona virus^1,2^. The first COVID-19 disease infected patient was identified in Wuhan, Hubei province, China on December, 2019. Till the reporting date, SARS-CoV-2 has infected 216 countries, area or territories and in total 4,731,458 cases and 316,169 deaths were confirmed^3,4^.

Variant types of comorbidities like diabetes, hypertension, cardiovascular disease are marked as the potential risk factors of COVID-19 disease severity^5^. However, Cancer patients are found as being more vulnerable to SARS-CoV-2 mediated infection rather than the persons having other disease complications because of their suppressive immunogenic state^6^. Recent investigation by a Chinese research team suggests, 18 out of 1590 cases of COVID-19 patients had a cancer history. Moreover, among these 18 patients, 5 patients had lung cancer (28%). In terms of fatality rate, a recent study in New York provides information that 6 out of 11 (55%) lung cancer patients died when infected by COVID-19^7,8^. All of these statistical reports remark the higher susceptibility and fatality of lung cancer patients due to COVID-19 infection. In this study, we aimed to find out the possible reasons behind this adverse outcome by targeting two crucial genes responsible for COVID-19 disease development, namely angiotensin-converting enzyme 2 (ACE2) and C-X-C motif chemokine 10 (CXCL10).

ACE2 acts as the receptor of host cells for the SARS CoV-2 viral spike-protein S1. For that reason, the entry of the viral particle is directly interconnected with the binding affinity of the ACE2 receptor^9,10^ On the other hand, CXCL10 is reported as a crucial cytokine plays significant role in proceeding the immune response to COVID-19 infected cells. In case of severe condition, the secretion of CXCL10 and other cytokines is exceedingly over-regulated that causes adverse assembly of cytokines which is known as cytokine storm (figure 1). This cytokine storm is responsible for the rapid development of acute respiratory distress syndrome (ARDS). In case of a prolonged severe condition, the cytokine storm also causes the alveolar collapse by destruction of the lung cells^11,12^. Concerning these adverse outcomes, we selected ACE2 and CXCL10 to identify the possible reasons for increased fatality rate of lung cancer patients in case of COVID-19. Overall, we aimed for a comprehensive in silico analysis on these two targeted genes to interpret the susceptibility and fatality of lung cancer patients towards COVID-19 infection by using available validated data of multidisciplinary databases.

**Figure 1.**
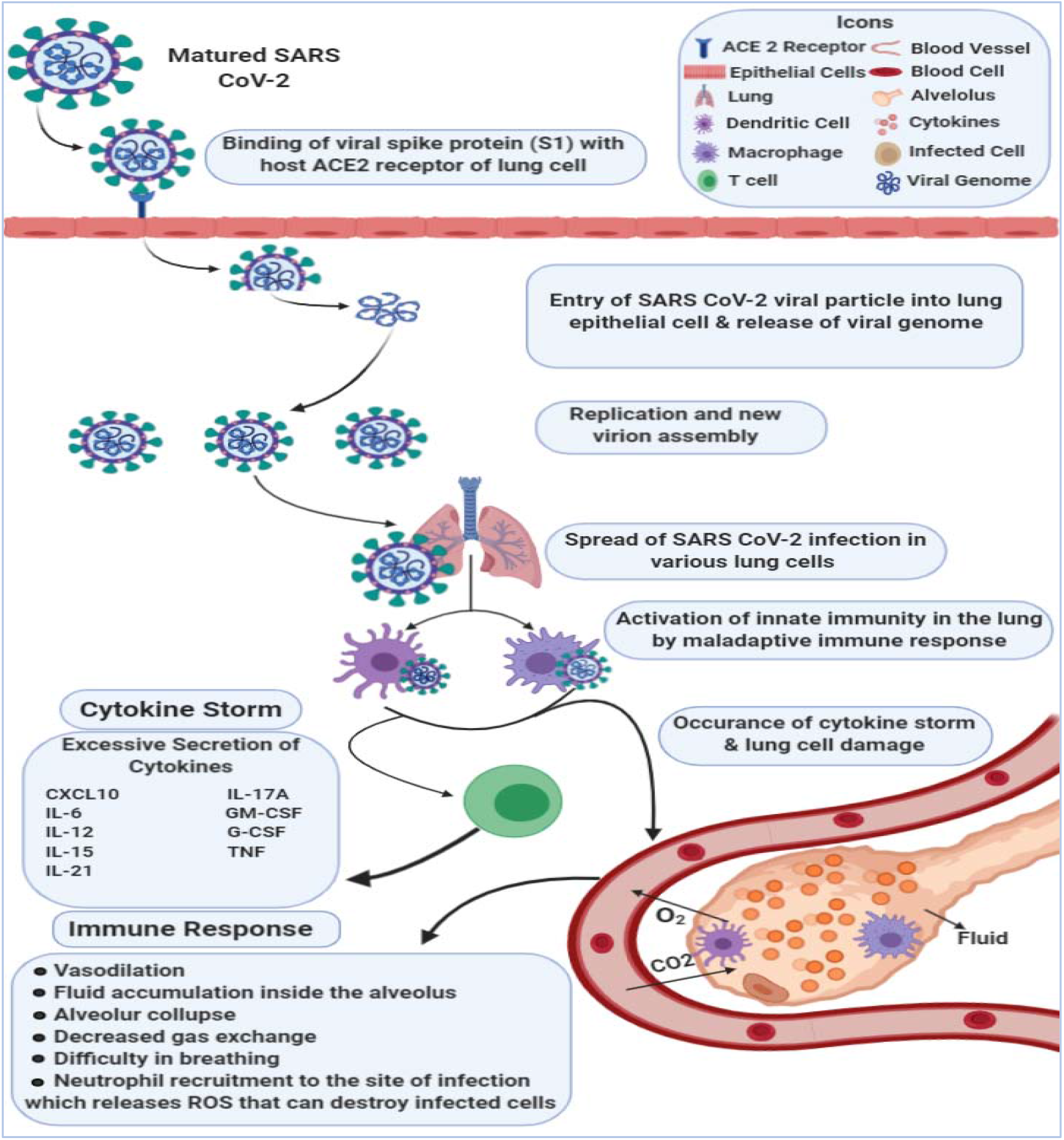
A Schematic diagram representing the cytokine storm assisted by the Cytokines.

## 2. Methods

### 2.1 Expression analysis of the targeted genes

#### 2.1.1 Analysis of gene expression in lung cancer

Data corresponding to ACE2 and CXCL10 mRNA expression pattern in lung cancer was retrieved from Tumor Immune Estimation Resource (TIMER) (http://cistrome.org/TIMER/). This tool allows comprehensive analysis of the immune suppressive nature of diverse cancer types. This web resource consists of six principal catalogs of analysis which allow to analyze the gene expression data and their interconnections along with the immune suppressed cancer cells^13^.

#### 2.1.2 Extensive Analysis of gene expression in LUAD and LUSC

In-depth expression analysis of ACE2 and CXCL10 in LUAD and LUSC was done by using the TCGA data from the UALCAN web portal (http://ualcan.path.uab.edu/). This is a public accessible web platform that is frequently used to analyze the expression, co-expression of multiple genes and their association with the clinical prognosis of variant cancer types by utilizing the TCGA data. UALCAN provides the analytical data for 31 types of cancer^14^.

#### 2.1.3 Correlation analysis of the targeted genes in LUAD and LUSC

Gene expression profiling interactive analysis (GEPIA) 2 database (http://gepia2.cancer-pku.cn/#index) is a web portal to analyze the mRNA expression of 8,587 normal and 9,736 tumor samples using the TCGA data. Besides the expression analysis, this database is also used for the gene specific correlation analysis of different cancer types^15^. Therefore, investigation of the correlation impact of ACE2 and CXCL10 in LUAD and LUSC was carried out by using the GEPIA 2 web portal.

### 2.2 Functional characterization of the targeted genes

#### 2.2.1 Mutation and CNAs determination in the targeted proteins

Exploration of significant genetic changes in ACE2 and CXCL10 was done by using cBioPortal (https://www.cbioportal.org/). This software is an interactive web portal for systematic analysis of the multidisciplinary data sets of cancer genomics. Providing the data from more than 5,000 tumor samples, this database is mostly used for the molecular profiling of cell lines and cancer tissue, mapping the frequency of mutations and other genetic alterations utilizing multidimensional cancer studies^16^.

#### 2.2.2 Development of protein-protein interaction network

Genes that are involved in COVID-19 development and predicted to have a dominant association with the viral disease were retrieved from the Comparative Toxicogenomics Database (CTD) (http://ctdbase.org/). CTD is a regularly updated database that delivers readily understandable records representing the gene-disease relationships^17^. To analyze the interconnection between the proteins significantly associated with COVID-19, a network was generated among them by using the STRING software (https://string-db.org/). This software is mainly used to construct protein-protein interaction network for showing multi-variant associations among the targeted proteins^18^.

### 2.3 Co-expression analysis of the targeted genes

#### 2.3.1 Identification of the co-expressed genes

The genes co-altered with the expression of ACE2 and CXCL10 in lung cancer were identified by using the R2: Genomics and Visualization platform (https://hgserver1.amc.nl/cgi-bin/r2/main.cgi). This database is an enriched resource of gene specific datasets that allows to analyze, interpret and reveal prominent outcomes of clinical research studies^19^.

#### 2.3.2 Determination of commonly co-expressed genes

The commonly co-altered genes of ACE2 and CXCL10 in both lung cancer and COVID-19 are determined by constructing the Venn diagrams using the Bioinformatics and Evolutionary Genomics (http://bioinformatics.psb.ugent.be/webtools/Venn/) web portal. The Venn diagram can generate a graphical output which represents the common elements of each listed intersection^20^.

### 2.4 Interpretation of functional role of the targeted genes

An integrative molecular assessment of the functional approaches of ACE2 and CXCL10 associated with the lung cancer and fatal type of COVID-19 development was attributed by using Protein Analysis Through Evolutionary Relationships (PANTHER) (http://www.pantherdb.org/) tool. This is a comprehensive tool that can classify genes in terms of various attributes such as biological procedures, cellular structures, molecular attitudes and pathways^21,22^.

## 3. Results

### 3.1 Expression of ACE2 and CXCL10 in lung cancer

Firstly, expression level of ACE2 and CXCL10 was analyzed by using TIMER web tool. Here, we found that ACE2 mRNA expression was upregulated in the lung adenocarcinoma (LUAD) and lung squamous cell carcinoma (LUSC) where p-value for LUAD was < 0.05 (Figure 3a). Increased level of expression was also showed by CXCL10 when compared to its normal tissues. P-value < 0.01 was evidenced in both the cancer types for CXCL10 (Figure 3b). Overall, these results gave a clear indication that ACE2 and CXCL10 are overexpressed in LUAD and LUSC.

**Figure 2.**
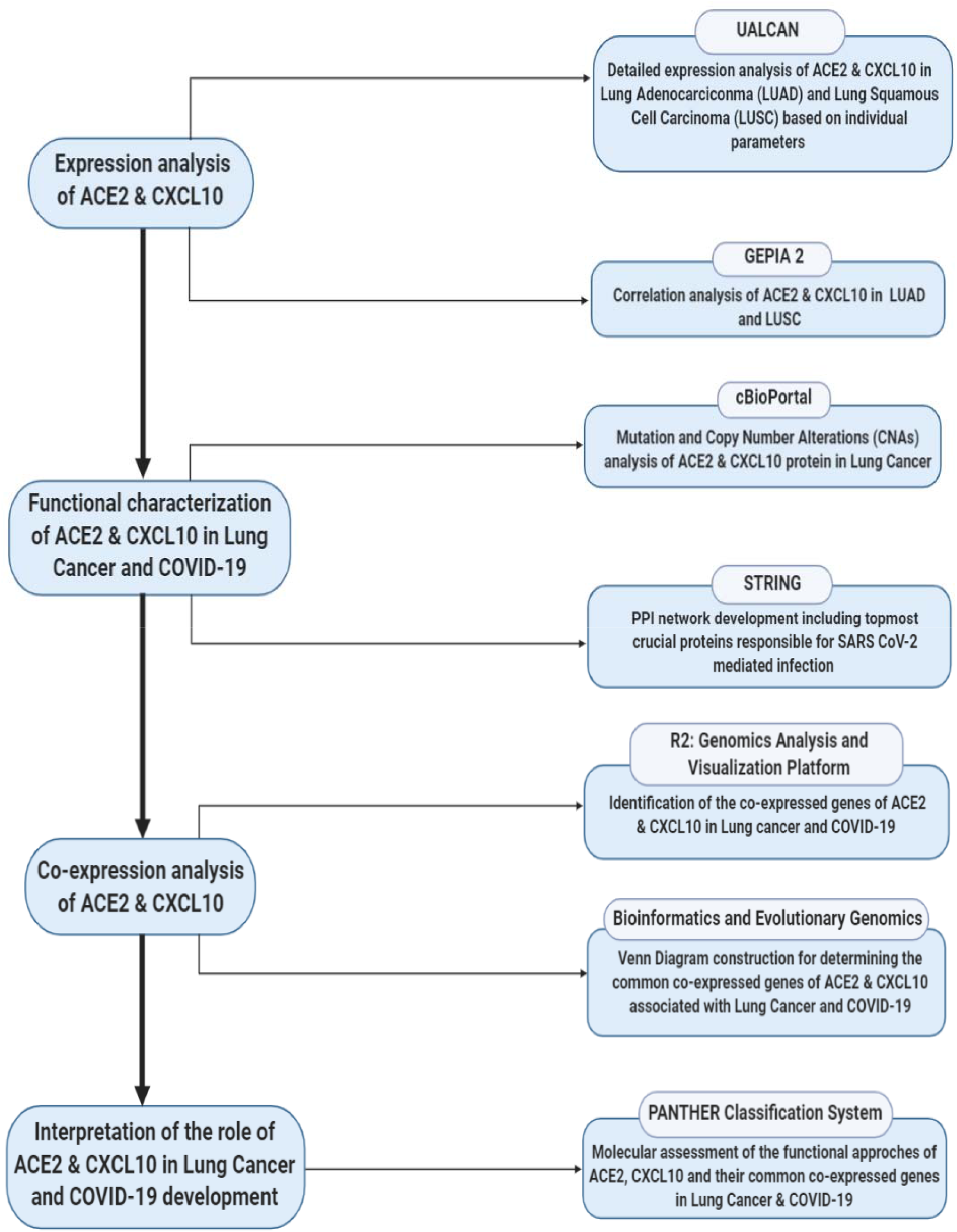
A schematic diagram representing the overall workflow of the study.

**Figure 3.**
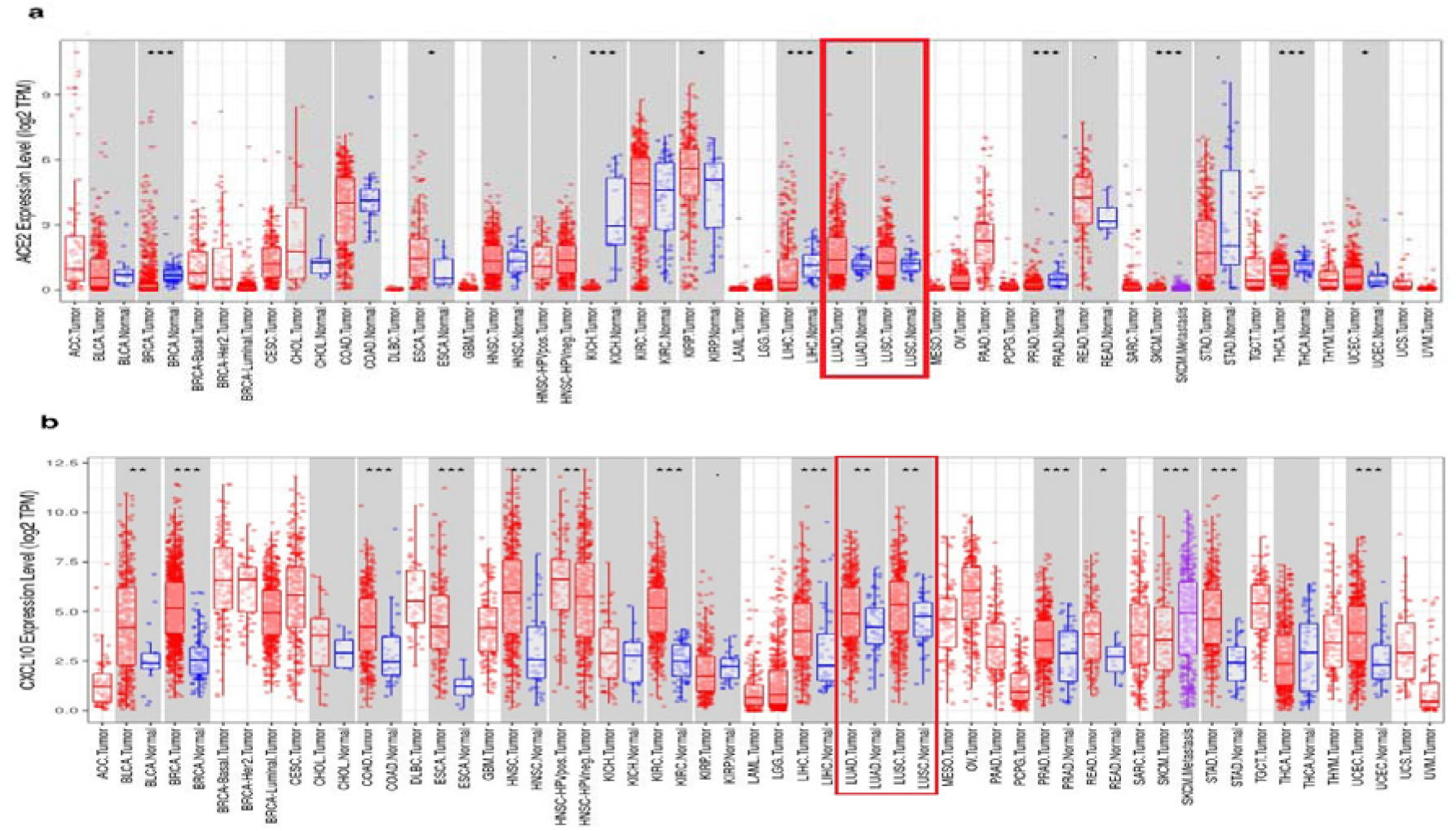
The expression analysis of ACE2 and CXCL10 by using TIMER (a) Different level of ACE2 mRNA expression is shown in multiple cancer studies whereas the overexpression of ACE2 in LUAD and LUSC is marked in red box (b) Different level of CXCL10 mRNA expression is shown in multiple cancer studies whereas the overexpression of CXCL10 in LUAD and LUSC is marked in red box. *P*-value codes: *<0.05, **<0.01, ***<0.001

### 3.2. Expression pattern and correlation analysis of ACE2 and CXCL10 in LUAD and LUSC

Using the UALCAN web tool, we found an upregulated expression pattern of ACE2 in LUAD and LUSC compared to the normal condition which is done on the basis of the TCGA dataset of individual cancer stages and different age groups (Fig 4a-d). In terms of individual cancer stages compared to normal condition, the ACE2 expression level was found higher in stage 1-3 but not in stage 4 in case of LUSC (Fig 4a-b). A similar kind of outcome representing the overexpression of ACE2 in terms of variant age groups was found, though for age group of 41-60 years, a surprisingly down-regulation of the gene expression was found (Fig 4c-d). On the other hand, for CXCL10 a significant over-expression was found in all cancer stages and age groups for both LUAD and LUSC patients (Fig 4e-h).

**Figure 4.**
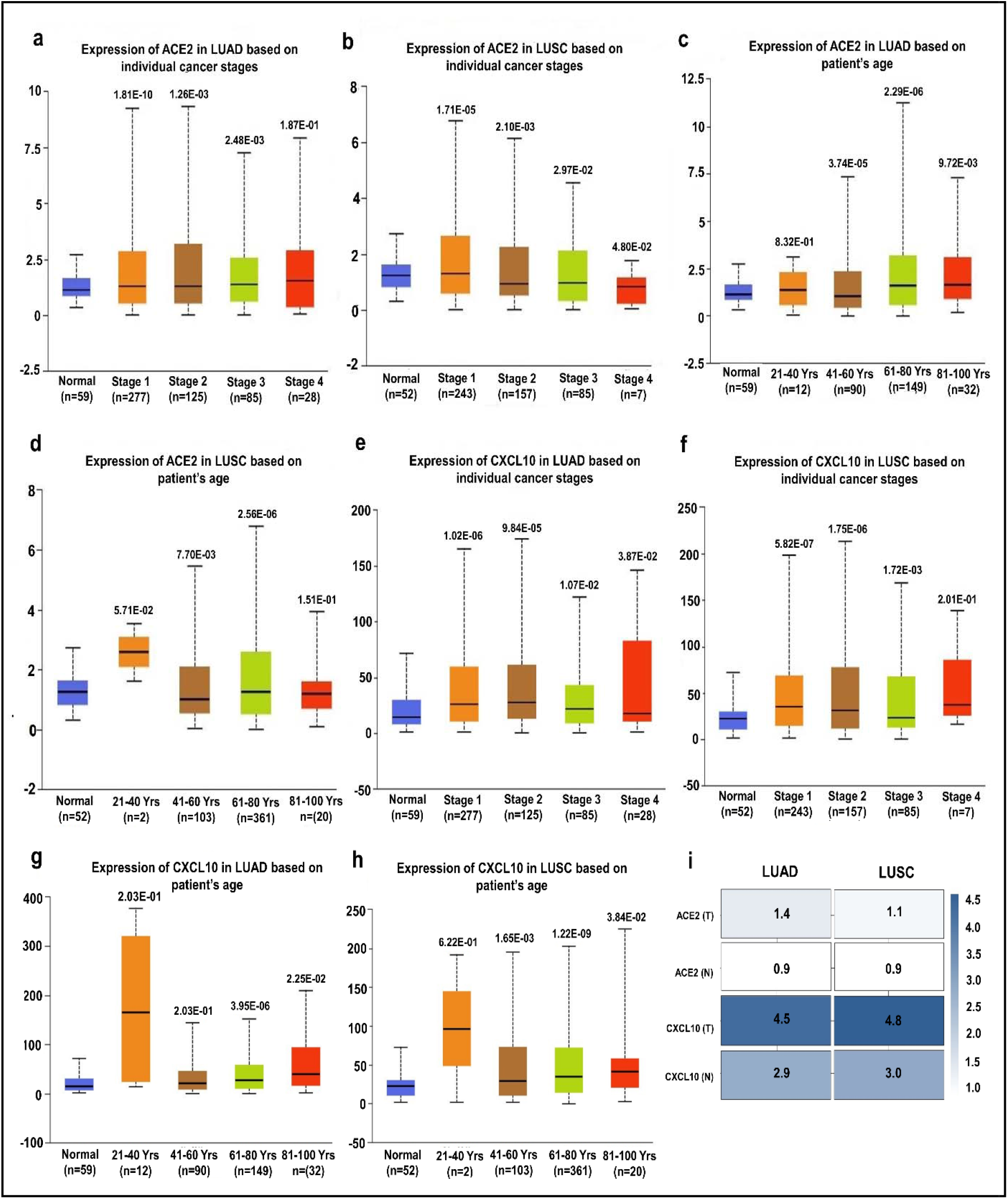
Expression analysis of ACE2 and CXCL10 in lung cancer using the UALCAN database (**a**) ACE2 gene expression in LUAD based on different cancer stages (**b**) ACE2 gene expression in LUSC based on different cancer stages; (**c)** ACE2 gene expression in LUAD based on different age groups(**d)** ACE2 gene expression in LUSC based on different age groups; (**e**) CXCL10 gene expression in LUAD based on different cancer stages (**f**) Expression level of CXCL10 gene in LUSC based on different cancer stages (**g**) CXCL10 gene expression in LUAD based on different age groups (**h**) CXCL10 gene expression in LUSC based on different age groups (**i**) Correlation analysis of ACE2 and CXCL10 gene expression in LUAD and LUSC based on tumor tissues (T) and normal lung tissues (N).

After that, using the GEPIA2 web portal, we generated an interactive heatmap which represents the correlation of ACE2 and CXCL10 along with both the cancer types, LUAD and LUSC on the basis of their expression score. Both the genes showed a deeper correlation when they were expressed by the tumor cells in comparison with normal cells (Fig 4i).

### 3.3. Analysis of genetic changes in ACE2 and CXCL10 protein sequences associated with lung cancer development

To evaluate the functional significance of ACE2 and CXCL10 in lung cancer development, we generated data representing multiple genetic alterations in ACE2 and CXCL10 mRNA using the cBioPortal database. Firstly, we prepared a query for ACE2 in this database using 6075 samples of 5719 lung cancer patients from 21 studies. From this analysis, we found out 64 mutations at 24 different locations of the 805 amino acids long human ACE2 protein. Out of these 64 mutations, 51 were missense type, while other 13 were recognized as truncated type (Fig 5a). After that, we extended our analysis to explore the frequency of genetic alterations in the ACE2 gene by using data from different lung cancer studies. Through this analysis, we observed that ACE2 is mostly altered in LUSC ranging the highest frequency of 3.49%. Though the alteration frequency fluctuated variously in multiple types of lung cancer studies, we managed to reveal that the least rate of alteration occurs in small cell lung cancer (figure 5b). Then, we focused on the expression level analysis of unique types of genetic alteration. In this case, we found that the highest level of mutation is occurred due to the shallow deletion type of genetic alteration. The second most significant genetic change is done by amplification type of copy number alteration. Variant natures of genetic alteration in ACE2 are assembled to contribute in the lung cancer development (Figure 5c).

**Figure 5.**
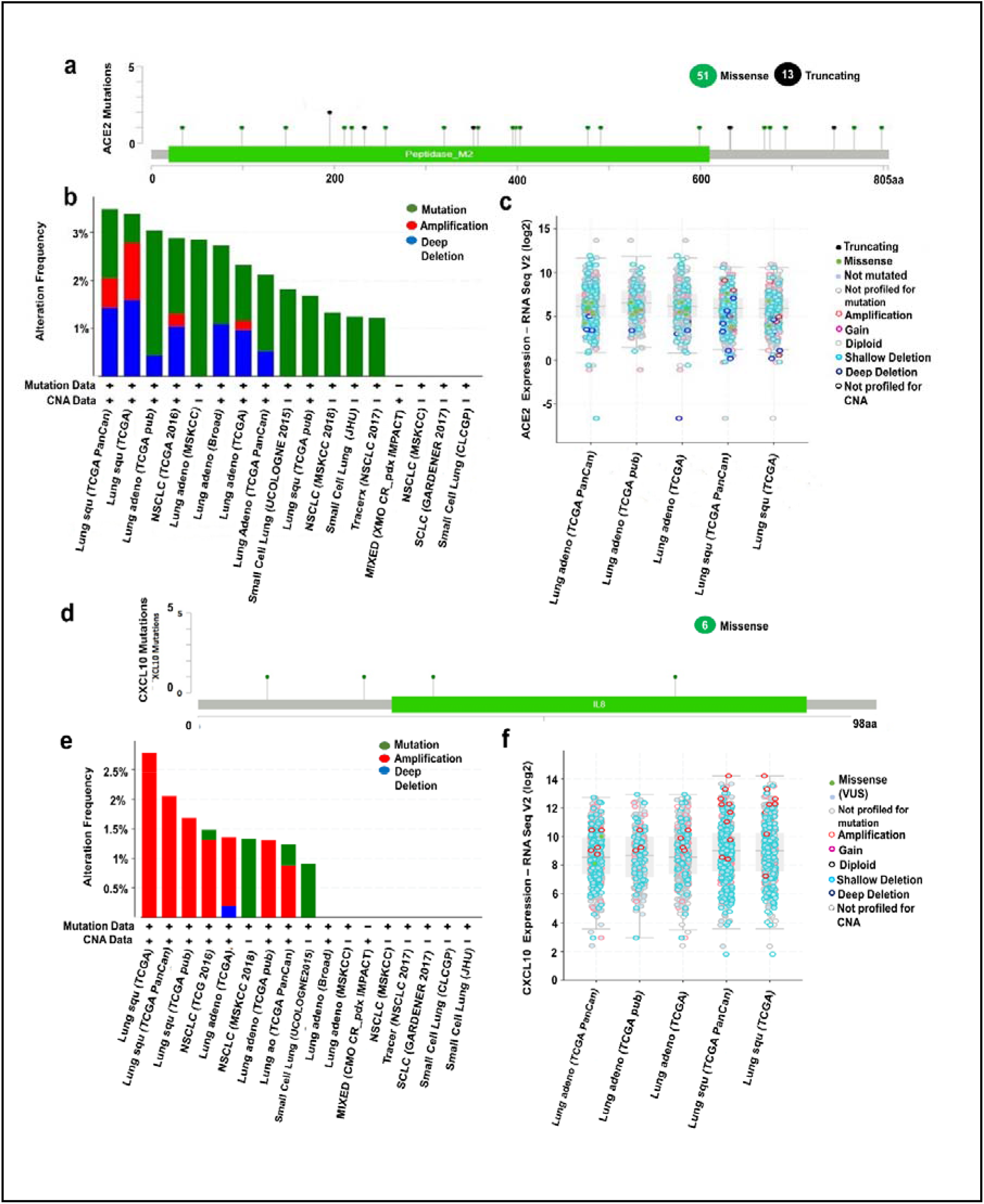
The Functional characterization of ACE2 and CXCL10 in lung cancer development by using the cBioPortal (**a**) 64 Mutations in ACE2 protein sequence was figured out by using lollipop plots (**b**) Three types of alteration frequencies of ACE2 in lung cancer were presented in bar diagram (**c**) The expression level of differently categorized genetic alterations was presented for ACE2 (**d**) Total 6 mutation in CXCL10 protein was presented by lollipop plots. (**e**) Three variant types of alteration frequency in CXCL10 were presented in the bar diagrams. (**f**) The expression level regarding to the multiple categories of genetic alteration was represented in graphical plots based on the log scale (RNA seqV2).

Next, we went through a similar type of analysis for interpreting the genetic alterations in the CXCL10 mRNA regarding lung cancer. In this instance, we pointed out 6 missense type of mutations with a somatic mutation frequency of 0.1% where 2 duplicate mutations were present. The mutations are evidenced at 4 different locations of the 98 amino acids long CXCL10 protein sequence (figure 5d). We then analyze genetic alteration frequency in the CXCL10 mRNA distinct numbers of lung cancer studies. In this regard, we observed the highest rate of alteration frequency (2.79%) in CXCL10 mRNA for LUSC. Interestingly, amplification is the one and only type of alteration in this case. However, for other cases, fluctuation in alteration frequency and changes in the nature of alteration was observed. For instance, the TCGA data of LUSC represented alteration frequency only for amplification type whereas UCOLOGENE 2015 revealed that alterations in CXCL10 of small cell lung cancer are completely mediated by mutation type genetic change (Figure 5e). Last of all, we did an extensive analysis of the different expression levels of unique alteration types in CXCL10 mRNA. From this analysis, we experienced that amplification is the most upregulated copy number alteration considering their level of expression while shallow deletion is the most common type of alteration considering their expression frequency (Figure 5f). Overall, the functional characterization of ACE2 and CXCL10 by using multiple lung cancer studies provides some of the vital evidence of their intimate relationship with lung cancer development.

### 3.4. Evaluation of the ACE2 and CXCL10 assisted protein-protein interaction network

Multiple numbers of proteins are responsible directly or indirectly for COVID-19 development. To explore the interconnection among the most vital proteins associated directly with this disease, first, we used the Comparative Toxicogenomics Database (CTD). Here, we managed to generate data of 12,347 genes associated with COVID-19 disease based on their inference score. Most importantly, each of the genes has either curated association to the disease or an inferred association via a curated chemical interaction. Extracting the huge data set, the number of 15 curated genes were identified as the biomarkers or therapeutic targets for COVID-19 treatment according to the information provided by CTD in which ACE2 and CXCL10 were included. By utilizing the translated protein sequences of these 15 genes a protein-protein interaction network was constructed through the STRING database (Figure 6). From this, we found out 74 connecting edges among 15 nodes of the selected proteins, though the expected edges were only 16 according to the information provided by the database itself. That means the network represents more interconnections than the expected outcome. Such affluence indicates that the proteins are functionally connected, as a group. Moreover, we found multiple numbers of interconnection for ACE2 and CXCL10 protein along with other topmost significant protein components associated with COVID-19 development. It is clear evidence that ACE2 and CXCL10 are crucial proteins contributing to the COVID-19 disease progression.

**Figure 6.**
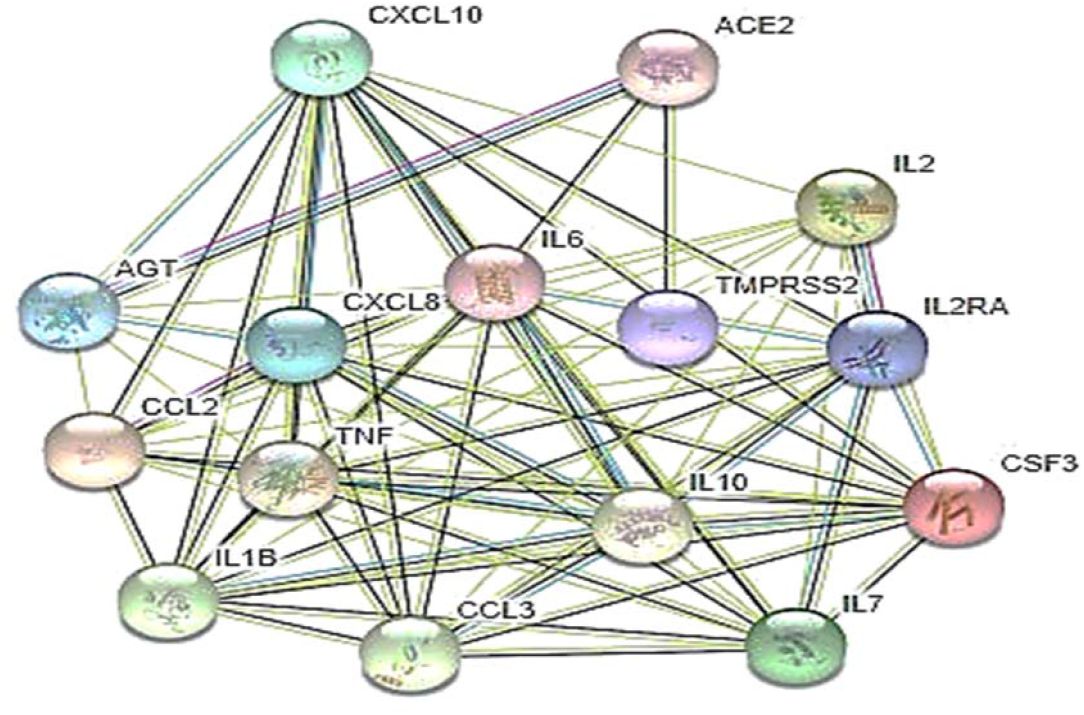
A protein-protein interaction network representing the interconnection of functional proteins associated with COVID-19 infection

### 3.5. Estimation of the commonly co-expressed genes of ACE2 and CXCL10 in lung cancer and COVID-19

To identify the genes that are correlated with the expression of ACE2 and CXCL10, we went through a comprehensive analysis by using the R2: Genomics and Visualization web portal. From here, we explored the co-expressed genes of ACE2 and CXCL10 responsible for lung cancer and COVID-19 development by utilizing the TCGA data. Here, we identified a total 6793 co-expressed genes of the ACE2 associated with LUAD and LUSC whereas the number of co-expressed genes related to COVID-19 was 10803. On the other hand, we determined 5999 genes that are co-altered with CXCL10 in case of lung cancer development and 6430 co-expressed genes associated with COVID-19. A restriction of *p*-value < 0.01 was applied to each case of analysis. After that, the lists representing co-expressed genes of ACE2 and CXCL10 in both cases of lung cancer and COVID-19 were utilized to construct Venn diagrams by using the Bioinformatics and Evolutionary Genomics web tool. The two Venn diagrams revealed the commonly co-expressed genes of ACE2 and CXCL10 associated with lung cancer and COVID-19 development. In case of ACE2, 3544 co-expressed genes were identified commonly associated with both of the disease conditions (Figure 7a). On the other hand, 2088 genes were identified which are co-altered along with the expression of CXCL10 in both cases of lung cancer and COVID-19 (Figure 7b).

**Figure 7.**
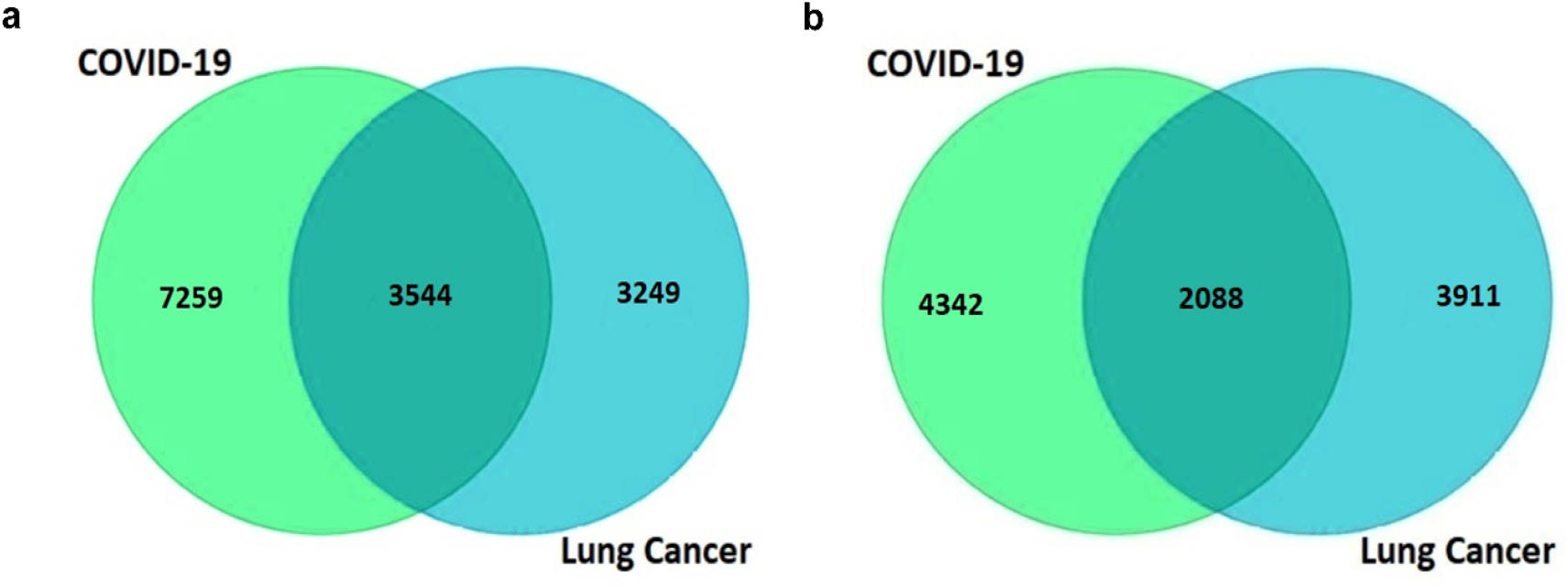
Graphical representation of commonly co-expressed genes of ACE2 and CXCL10 in lung cancer and COVID-19 **(a)** The Venn diagram represents 3544 co-expressed genes of ACE2 **(b)** The Venn diagram represents 2088 commonly co-expressed genes of CXCL10 associated with both of the diseases.

### 3.6. Molecular assessment of the functional approaches of ACE2 and CXCL10

To interpret the functional activity of ACE2 and CXCL10 in lung cancer and COVID-19, we used a list of genes including the targeted ones and their commonly co-expressed genes associated with both of these diseases using the PANTHER data analysis platform. First of all, we processed a query for determining the molecular activity of ACE2 by listing the previously identified 3544 commonly co-expressed genes. Following the next steps, we analyzed multiple types of molecular activity in which a major portion (37.1%) of the genes (803) are involved in the binding activity (Figure 8a). Through extended analysis, we observed the variant nature of the binding activities of corresponding 803 genes. In this instance, we found out that most of the genes are involved in protein binding activity (53.5%; 430 genes) (Figure 8b).

**Figure 8.**
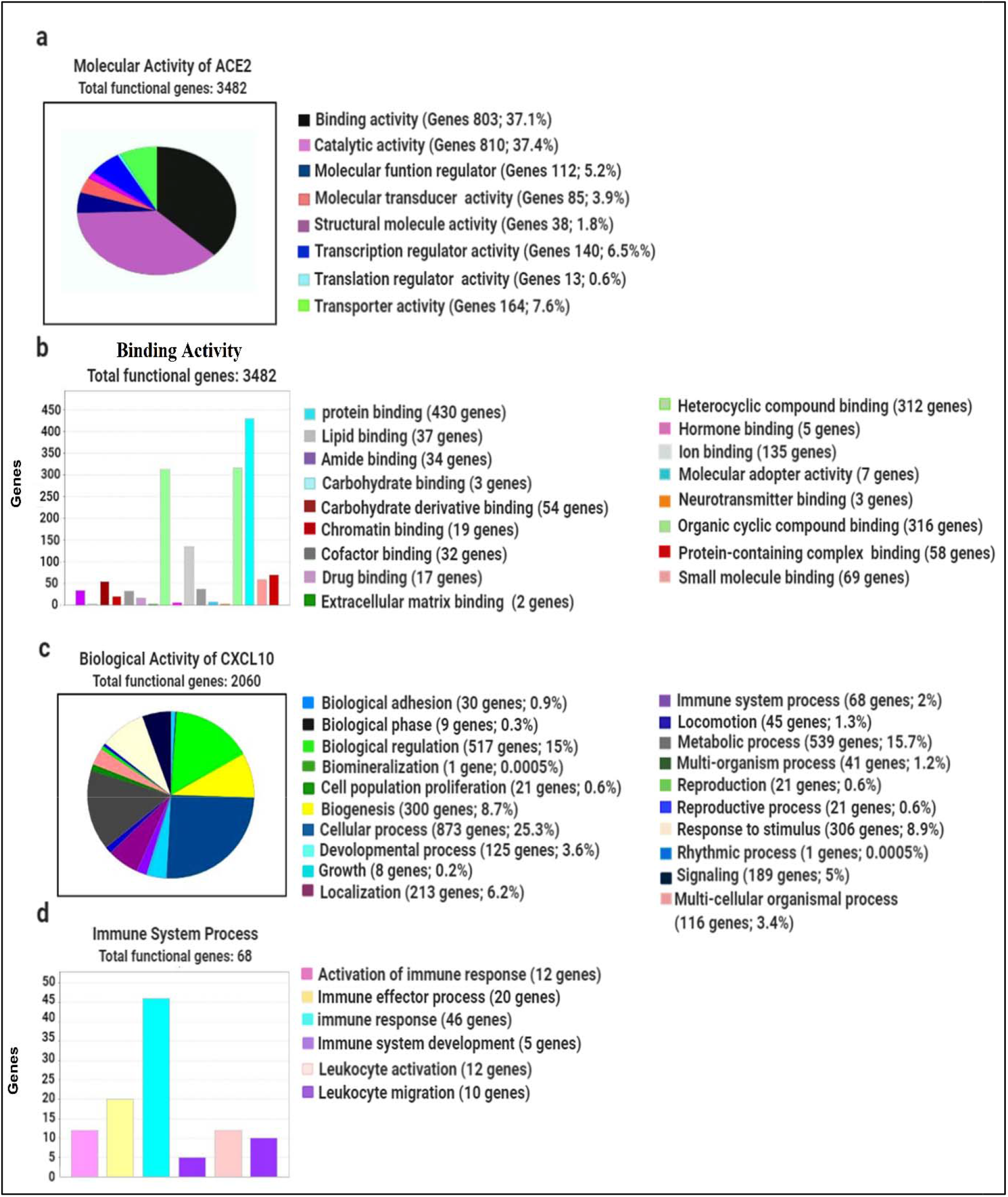
Evaluation of the functional attitudes of ACE2 and CXCL10 represented by using the PANTHER database **(a)** Eight classes of the molecular activity of ACE2 and its coexpressed genes were presented through a pie chart **(b)** In total 17 variant types of binding activities of 803 genes were represented by using a bar chart **(c)** Total 20 unique types of biological activities of CXCL10 and its co-expressed genes were represented using a pie chart **(d)** Six differently categorized immune system processes of the corresponding 68 genes were presented through a bar chart.

After that, we looked forward to investigating the functional attitude of CXCL10 using the list of previously determined 2048 commonly co-expressed genes associated with lung cancer and COVID-19 development. By analyzing the biological processes, we observed that the listed genes are involved in performing a wide range of biological activities such as cellular process, biological regulation, metabolic process, immune system process, localization, biogenesis etc (Figure 8c). However, by remarking our research aims, we further looked forward only to do an extended analysis with the genes involved in different immune processes. Regarding this analysis, we explored that 46 genes are involved in immune response, 5 genes act for immune system development, 12 genes are responsible for leukocyte activation whereas 10 genes are responsible for leukocyte migration, 12 genes contribute to the activation of immune response and 20 genes lead the immune effector process (Figure 8d). Overall, along with the functional activity of CXCL10, many of the coexpressed genes are found to play important in various branches of the immune system.

## 4. Discussion

Recently, COVID-19 is remarked as the most threatened clinical pandemic of the century which is showing more vulnerable attitudes towards cancer patients. Current statistical reports claimed that among these COVID-19 infected cancer patients, the most frequent and severe nature of infection has been found against lung cancer^7,23^.

In this research study, we aimed to find out the possible reasons behind the higher susceptibility and fatality rate of lung cancer patients to SARS CoV-2 mediated infection. To proceed with an impactful study, we targeted the expression analysis of ACE2 and CXCL10 because of their crucial roles in viral entry and disease severity, respectively. The newest scientific outcomes revealed that ACE2 acts as the main receptor of SARS CoV-2. Moreover, the susceptibility of COVID-19 patients is directly interconnected with the binding affinity of ACE2 to the S1 spike protein which is known as the receptor-binding domain of the virus^24^. On the other hand, CXCL10 is treated as one of the significant cytokines which is directly involved in the occurrence of cytokine storm at the severe condition of the COVID-19 infection. This cytokine storm is directly associated with the prosecution of Acute Respiratory Disease Syndrome (ARDS)^11,25^. Moreover, at the severe stage of infection, the cytokine storm causes the excessive secretion of cytokines in the alveoli. This enormous assembly of cytokines suddenly gets involved in a suicidal approach by causing the damage of the lung cells instead of providing the required immune response^26,27^. For being involved with this severe prognosis of COVID-19, CXCL10 is predicted as a significant factor for the fatality of COVID-19 patients. On the basis of these evidence, we analyzed the level of ACE2 mRNA expression in LUAD and LUSC attributing multidisciplinary parameters. Interestingly almost in all of the cases, we found an over-expression of ACE2 and CXCL10 in LUAD and LUSC compared to the normal condition. However, two contradictory results of ACE2 expression were found in individual cancer stage-4 of LUSC and age group 41-60 years of LUAD and LUSC. Additionally, we applied a correlation analysis of ACE2 and CXCL10 with LUAD and LUSC on the basis of their expression score by normal and tumor cells. In this case, we observed a much deeper correlation of these two genes with LUAD and LUSC when they are expressed by tumor cells. The results of this initial analysis may provide a primary evidence that overexpression of ACE2 receptor and CXCL10 cytokine is one of the major possible reasons behind increased susceptibility and fatality of lung cancer patients towards COVID-19. To make the evidence stronger enough, we extended the analytical process to the investigation on the functional assessment of ACE2 and CXCL10.

From the updated scientific outcomes, it is already clear that ACE2 and CXCL10 have a strong correlation with COVID-19 infection development^2,11^. However, in the next part of this study, we intended to provide an additional evidence to this scientific knowledge by establishing a protein-protein interaction network among the topmost significant 15 proteins responsible for the COVID-19 development including ACE2 and CXCL10. Following this analysis, we found 74 nodes of interaction among the 15 proteins where the major portion of the nodes based protein network is assisted by ACE2 and CXCL10. This provides an extended evidence about the functional involvement and significant contribution of ACE2 and CXCL10 to the SARS CoV-2 mediated infection development. Besides confirming the role of ACE2 and CXCL10 in COVID-19 development, we also realized the need of determining their role in case of lung cancer development to establish a connection between lung cancer and COVID-19 through these targeted genes. Therefore, we did the functional characterization of ACE2 and CXCL10 in lung cancer by analyzing the mutations and copy number alterations in their protein sequence based on 21 lung cancer studies. In total 64 mutations at 24 different locations of the ACE2 protein were found where the maximum frequency of alteration is 3.49 %. On the other hand, 6 mutations at 4 different locations of the protein sequence were found against CXCL10 where the highest level of alteration frequency was 2.79%. For both cases, we observed that shallow deletion is the most frequent type of CNAs. These results are supportive evidence assuring the active participation of ACE2 and CXCL10 in lung cancer development.

After that, we looked forward to identify the commonly co-expressed genes of ACE2 and CXCL10 associated with lung cancer and COVID-19. We constructed two Venn diagrams by listing the co-expressed genes of ACE2 and CXCL10 in each cases of lung cancer and COVID-19. Therefore, 3544 and 2088 commonly co-expressed genes associated with both lung cancer and COVID-19 were identified for ACE2 and CXCL10, respectively. These commonly co-expressed genes were utilized to interpret the functional attitudes of ACE2 and CXCL10 both in lung cancer and COVID-19. By analyzing the molecular activity of the 3544 commonly co-expressed genes for ACE2 we found that 803 genes including the ACE2 are involved in the binding activity. The active participation of these commonly co-expressed genes in binding activity can be beneficial for ACE2 to show a more efficient binding affinity towards SARS CoV-2 viral spike protein. This functional enforcement of the receptor activity of ACE2 may be increase the susceptibility of the lung cancer patients towards COVID-19. On the other hand, to interpret the functional attitude of CXCL10 we used the list of 2088 commonly co-expressed genes associated with lung cancer and COVID-19. By analyzing the biological activity of 2088 co-expressed genes, we found that 68 genes are directly involved in the immune system including the CXCL10. At the early stage of our study, we showed an over-expression of CXCL10 in lung cancer and by doing the functional analysis of CXCL10 we found its association with immune response. By combining these two analyses, we can say that over expression of CXCL10 along with the presence of other cytokines causes cytokine storm in the alveoli of the COVID-19 infected lung cancer patients. This cytokine storm results in the alveolar collapse due to lung cell damage by excessive neutrophil recruitment to the site of infection. In the prolonged severe condition, the damaged alveolus causes decreased gas exchange and results in difficulty in breathing. Finally, the ultimate destruction of the alveoli results in the stoppage of breathing and causes patient’s death^11,27^. Overall, enforcing the viral entry and cytokine storm, the over-expressed ACE2 and CXCL10 become major factors of the higher susceptibility and fatality rate of the lung cancer patients towards COVID-19 infection.

### Concluding Remarks

From this study, we have gained valuable insights suggesting that ACE2 and CXCL10 are vitally involved in SARS CoV-2 entry and its severity. The key findings of this research also represented the overexpression and direct interconnection of ACE2 and CXCL10 with lung cancer development. Therefore, on the basis of this systemic analysis, we can conclude that ACE2 and CXCL10 are possible biomarkers whose higher expressions are responsible for the greater susceptibility and fatality of lung cancer patients towards the COVID-19 infection. However, we encourage further wet-lab research on our study outcomes to generate extended information.

## Data Availability Statement

No data are associated with this article.

## Competing Interests

No competing interests were disclosed.

## Notes

### Competing Interest Statement

The authors have declared no competing interest.

